# An immune-suppressing protein in human endogenous retroviruses

**DOI:** 10.1101/2022.11.04.515135

**Authors:** Huan Zhang, Shengliang Ni, Martin C. Frith

**Affiliations:** Graduate School of Frontier Sciences, University of Tokyo, Chiba, Japan; Artificial Intelligence Research Center, AIST, Tokyo, Japan; Computational Bio Big-Data Open Innovation Laboratory, AIST, Tokyo, Japan

**Author notes:** Joint first authors (Huan Zhang, Shengliang Ni).

## Abstract

Retroviruses are important contributors to disease and evolution in vertebrates. Sometimes, retrovirus DNA is heritably inserted in a vertebrate genome: an endogenous retrovirus (ERV). Vertebrate genomes have many such virus-derived fragments, usually with mutations disabling their original functions.

Some primate ERVs appear to encode an overlooked protein. This protein is homologous to protein MC132 from Molluscum contagiosum virus, which is a human poxvirus, not a retrovirus. MC132 suppresses the immune system by targeting NF-κB, and it had no known homologs until now. The ERV homologs of MC132 in the human genome are mostly disrupted by mutations, but there is an intact copy on chromosome 4. We found homologs of MC132 in ERVs of apes, monkeys, and bushbaby, but not tarsiers, lemurs or non-primates. This suggests that some primate retroviruses had, or have, an extra immune-suppressing protein, which underwent horizontal genetic transfer between unrelated viruses.

## INTRODUCTION

Retroviruses cause significant disease, such as AIDS and adult T-cell leukemia. They have RNA genomes, which undergo reverse-transcription into DNA, which is inserted into the host cell’s genome. Occasionally, they infect germ-line cells, in which case the insertion may be inherited by future generations of the host organism: this is termed an endogenous retrovirus (ERV). Vertebrate genomes contain many retrovirus-derived fragments, for example they comprise ~8% of the human genome. Probably most ERVs decay by neutral evolution, however, many ERV fragments have been co-opted by the host to function as protein-coding genes or regulatory elements [1,2,3]. Thus, retroviruses are important contributors to disease and evolution.

Retroviruses encode three main genes, in the following order: 5’-gag-pol-env-3’. Each gene produces several proteins, including viral structural proteins, a protease, reverse transcriptase, and viral envelope proteins. Some retroviruses also encode small “accessory” proteins near the 3’-end. In short, retroviruses have a largely consistent genome organization.

We report that some primate ERVs encode an extra protein upstream of the gag gene (or possibly fused to the gag gene). This protein is homologous to protein MC132 of the human poxvirus Molluscum contagiosum. MC132 suppresses the immune system by targeting NF-κB, and it had no known homologs until now [4].

## RESULTS

### MC132 protein fossils in human ERVs

We discovered this ERV protein by chance, while searching for protein fossils in the human genome. Protein fossils are segments of formerly protein-coding DNA, and we recently developed a sensitive method to find them by comparing DNA to protein sequences to find homologous segments [5,6]. We found a few dozen segments of the human genome with homology to protein MC132 from Molluscum contagiosum virus (specifically, proteins Q98298 and A0A1S7DLX6 from UniProt release 2022_03). These segments lie in ERVs. There are many types of human ERV, and these segments lie in a few specific types. We used ERV annotation from RepeatMasker, which finds ERVs by comparing the genome to ERV models in the Dfam database [7]. These models and annotation are not perfect (see below), but according to them HERV30 has a large segment with high similarity to the protein (Figure 1), and there are weaker matches in HERV17, HERV9, and perhaps a few others. These ERV types are closely related: they are in Dfam’s ERV1 class.

**Figure 1:**
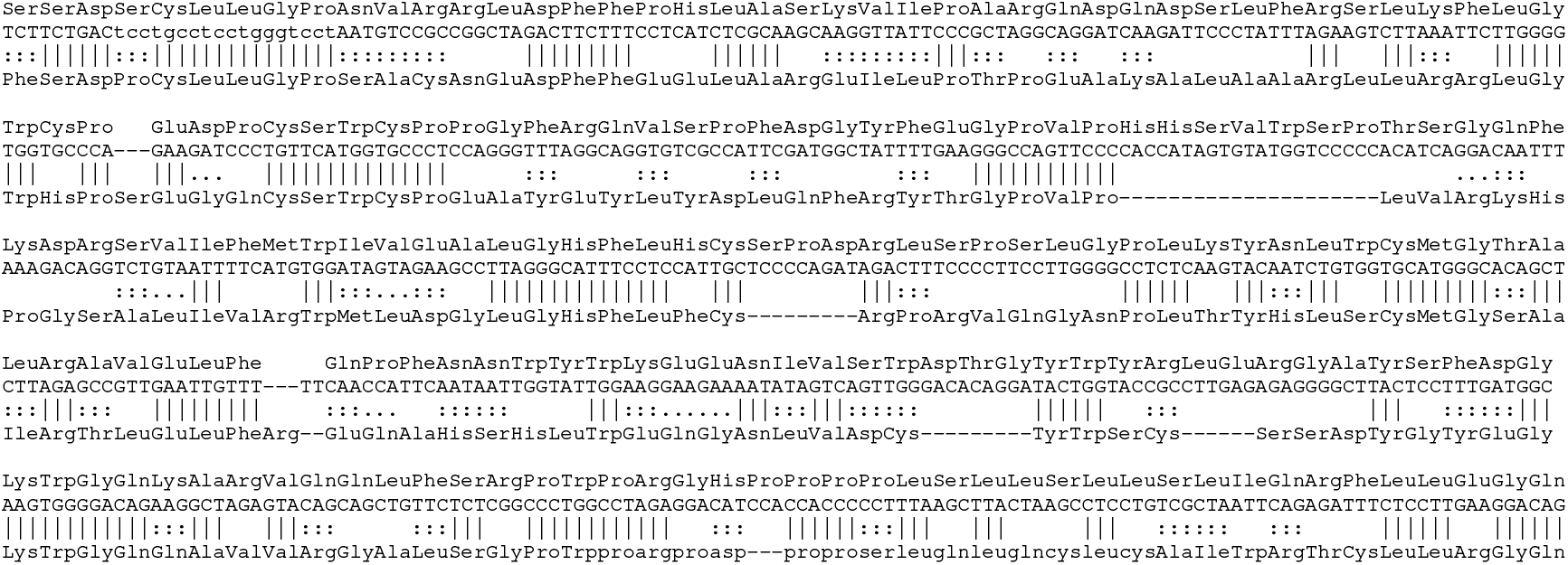
Alignment between Dfam’s HERV30 consensus DNA sequence and MC132 protein (Q98298). The DNA’s translation is shown above it. ||| indicates a match, ::: a positive substitution score, and … a zero substitution score. Lowercase regions were deemed to be simple repeats by the alignment tool (LAST).

### Reconstructing the ancestral sequence

These ERVs presumably inserted into the genome millions of years ago and underwent random mutations, degrading the original sequence. We attempted to reconstruct an ancestral protein, by taking 15 DNA segments with strong similarity to MC132, plus 200 bp flanks, and feeding them to Refiner [8]. Refiner inferred an ancestral DNA sequence, which remarkably has an 879 bp open reading frame (ORF) encompassing the MC132 homology. This ORF encodes the following protein:

`MAPPEAPPAXVTERETATSSDPCLLGPNVRRLDFFPHLASKVIPARQDQD`

SLFRSLKFLGWRPEDPCSWCPPGFRQVSPFDGYFEGPVPHHSVWSPTSGQ

FKDRSVIFMWIVEALGHFLHCSPDRLSPSLGPLKYNLWCMGTALRAVELL

FQPFNNWYWKEENIVSWDTGYWYRLERGAYSFDGKWGQKARVQQLFSRPW

PRGHPPPPLSLLSLLSLIQRFLLEGQFYGQAHVNWALACKHQWCPRPRPC

HPGTGRTRWQKDHNKSNSPCAPFSGQWAHGRGKGSFHPAGKHG

It is easy to verify (e.g. by NCBI BLAST) that this protein has significant similarity to MC132. The DNA from Refiner is 99% identical to Dfam’s HERV30 consensus sequence, but the latter has two frame-shifts disrupting the ORF.

We then sought human genome segments homologous to this new protein, and found hundreds of hits (Table 1), mostly in HERV17 annotations (Table 2). The hits are consistently upstream of the gag gene (Figure 2). Remarkably, there is one HERV30 in chromosome 4 where the ORF is intact, with no frame disruptions (Figure 2).

**Table 1:** Alignments of the reconstructed protein sequence and Q98298 in each genome

| Organism |  | Alignments |  |  |
| --- | --- | --- | --- | --- |
|  |  | Refiner sequence | Q98298 | Refiner+Q98298 |
| Human | *Homo sapiens* | 252 | 65 | 317 |
| Gibbon | *Nomascus leucogenys* | 221 | 46 | 267 |
| Rhesus | *Macaca mulatta* | 232 | 61 | 293 |
| Golden snub-nosed monkey | *Rhinopithecus roxellana* | 253 | 43 | 296 |
| Marmoset | *Callithrix jacchus* | 267 | 92 | 359 |
| Bushbaby | *Otolemur garnettii* | 7 | 4 | 11 |

**Table 2:** Overlaps between the protein matches and RepeatMasker ERV annotations

| Organism | ERV Overlaps between alignment and RepeatMasker |  |  |  |  |  |  |
| --- | --- | --- | --- | --- | --- | --- | --- |
|  | HERV17 | HERV9 | HERV9N | HERV30 | HERVIP10FH | HERVK14 | Other |
| Human | 156 | 45 | 39 | 23 | 8 | 2 | 33 |
| Gibbon | 120 | 45 | 33 | 16 | 5 | 3 | 9 |
| Rhesus | 155 | 100 | 0 | 18 | 5 | 2 | 4 |
| Golden snub-nosed monkey | 151 | 100 | 0 | 24 | 7 | 2 | 4 |

**Table 3:** Genome Versions

| Organism |  | Genome Assembly | RepeatMasker Version |
| --- | --- | --- | --- |
| Human | *Homo sapiens* | GRCh38.p14 | 4.1.0 |
| Gibbon | *Nomascus leucogenys* | Asia_NLE_v1 | 4.0.8 |
| Rhesus | *Macaca mulatta* | Mmul_10 | 4.0.8 |
| Golden snub-nosed monkey | *Rhinopithecus roxellana* | Novogene Rrox_v1 | 4.0.8 |
| Marmoset | *Callithrix jacchus* | Callithrix_jacchus_cj1700_1.1 | 4.0.8 |
| Tarsier | *Tarsius syrichta* | tarSyr2 | 4.0.5 |
| Mouse lemur | *Microcebus murinus* | Mmur_3.0 | 4.0.6 |
| Bushbaby | *Otolemur garnettii* | OtoGar3 | 4.0.6 |
| Norway rat | *Rattus norvegicus* | mRatBN7.2 | 4.0.8 |
| Pika | *Ochotona princeps* | OcjPri4.0 | 4.0.8 |
| Dolphin | *Tursiops truncatus* | mTurTru1.mat.Y | 4.0.8 |

**Figure 2:**
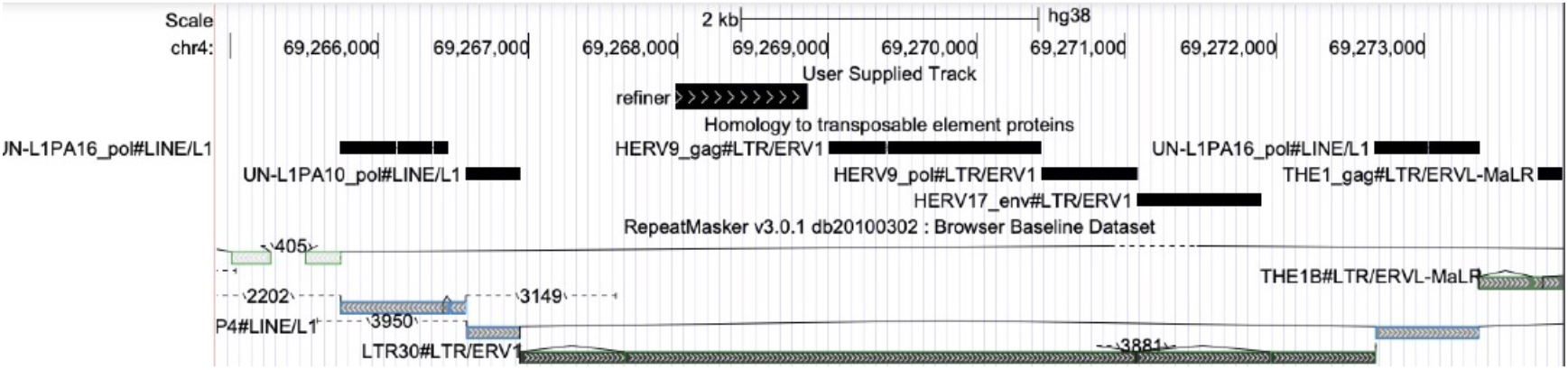
Location of the newly-discovered protein in an ERV in human chromosome 4. The location of the new protein is shown by the top bar labeled “refiner”. The black bars below that show DNA segments aligned to known transposable element proteins (from a previous study [6]), which are in the usual gag-pol-env order. Below that are RepeatMasker annotations of transposable element-derived segments. Here, RepeatMasker annotates two long terminal repeats of type LTR30, flanking an internal retroviral sequence of type HERV30. There is a 3881 bp deletion near the end of this internal sequence. Screenshot from the UCSC genome browser (http://genome.ucsc.edu) [9].

### The protein homology overlaps gaps in HERV17 annotations

We noticed that, in HERV17, the DNA region homologous to the new protein overlaps a consistent gap in RepeatMasker’s HERV17 annotation (Figure 3). This gap indicates an imperfection in the ERV models used by RepeatMasker. Either the HERV17 model is inaccurate in this region, or we have a new HERV17-like subfamily that is not yet represented in RepeatMasker’s models. We made a new model, by feeding HERV17 sequences from the human genome to Refiner. The new consensus sequence is 99% identical to Dfam’s HERV17 over most of its length, but has an extra 270 bp in the gap region. This suggests we should update the model rather than add a new subfamily. On the other hand, a previous study suggested two HERV17 subgroups: the extra 270 bp was not discussed, but is actually present in one subgroup [10]. In any case, the gap region is immediately upstream of a GA tandem repeat, which may evolve quickly and cause variation in these models.

**Figure 3:**
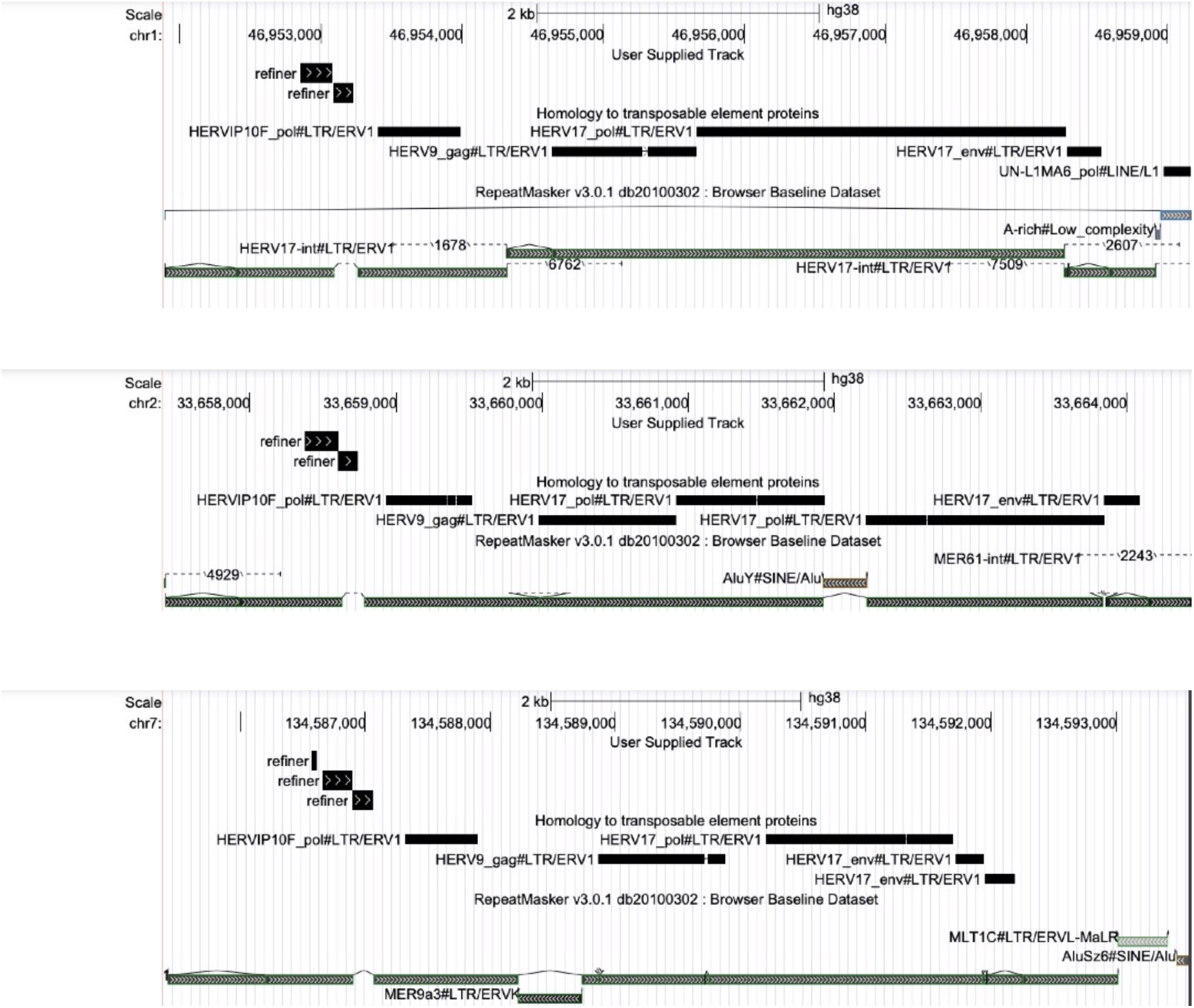
The new protein overlaps gaps in HERV17 annotations. The panels show 3 human genome locations with homology to the new protein (bars labeled “refiner”). The protein aligns to each location as 2 or 3 separate fragments. The black bars below that show DNA segments aligned to known transposable element proteins. Below that are RepeatMasker annotations of transposable element-derived segments. In each case, the new protein overlaps a retroviral sequence of type HERV17. However, in each case the protein overlaps a consistent gap in RepeatMasker’s HERV17 annotation. There is also an unexpected pol protein homology (HERVIP10F_pol) between the new protein and the gag gene.

Our new consensus sequence has frameshifts in the region homologous to the new protein. So do the previous subgroups. Thus, HERV17 may have proliferated in the genome after disruption of the reading frame. HERV17 (also known as HERV-W) is interesting because it has many copies that were retrotransposed by LINE enzymes [11], and its env gene was co-opted as the human syncytin gene *ERVW-1* involved in placental development [12].

### Extent of protein homology in ERV families

To better understand this protein homology in each ERV family, we took the genome segments aligned to the Refiner protein, and mapped them to Dfam’s consensus DNA sequence for each ERV (Figure 4). HERV17 consistently has a partial match, shorter than the HERV30 match, while HERV9 and HERV9N have even shorter matches.

**Figure 4:**
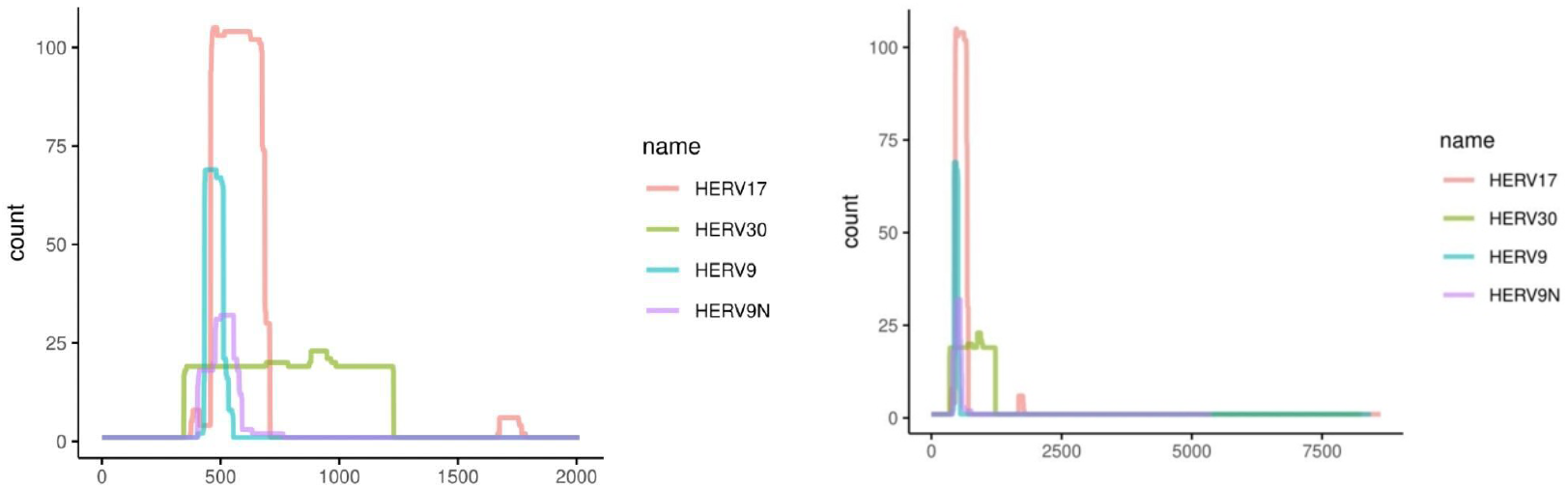
Locations of the newly-found protein in four ERV families. The right-hand panel shows the full-length ERVs, and the left-hand panel is zoomed in to the matching region. HERV30 has a long match to the protein (green), while the other ERVs have fragmentary matches. The separated matches in HERV17 (red) appear to correspond to the annotation gap shown in Figure 3.

In some of these cases, the RepeatMasker ERV annotations are fragmented and suggest ambiguity about which type of ERV1 is really present. It is possible that some of the Dfam consensus sequences incorrectly combine different ERV subfamilies, or that some ERVs are actually chimeric. Careful reconstruction of these ERVs would help us to understand the evolution of this protein.

### The new ERV protein in non-human primates

Finally, we searched several mammal genomes for homologies to MC132 and the new protein from Refiner. Similarly to human, there are hundreds of hits in apes, old-world monkeys, and new-world monkeys (Table 1). Among other primates, there are a few hits in bushbaby, but none in tarsier or mouse lemur. This is surprising, because it is usually thought that bushbaby and lemurs are related as Strepsirrhini, whereas tarsiers and simians are related as Haplorhini. We found no hits in other mammals (e.g. rat, pika, dolphin).

As expected, the homologous segments of these primate genomes lie in ERVs. In apes and old-world monkeys, these are the same types of ERV as in human, according to RepeatMasker annotations. In a more-distantly related new-world monkey (marmoset), we find the highest number of homologous segments, which almost all lie in ERV annotations of type ERV1-1_CJa-I (also in the ERV1 category). In bushbaby, the hits overlap gaps in ERV annotations (Figure 5). It is likely that the ERV models used by RepeatMasker become less accurate for primates more distantly related to human.

**Figure 5:**
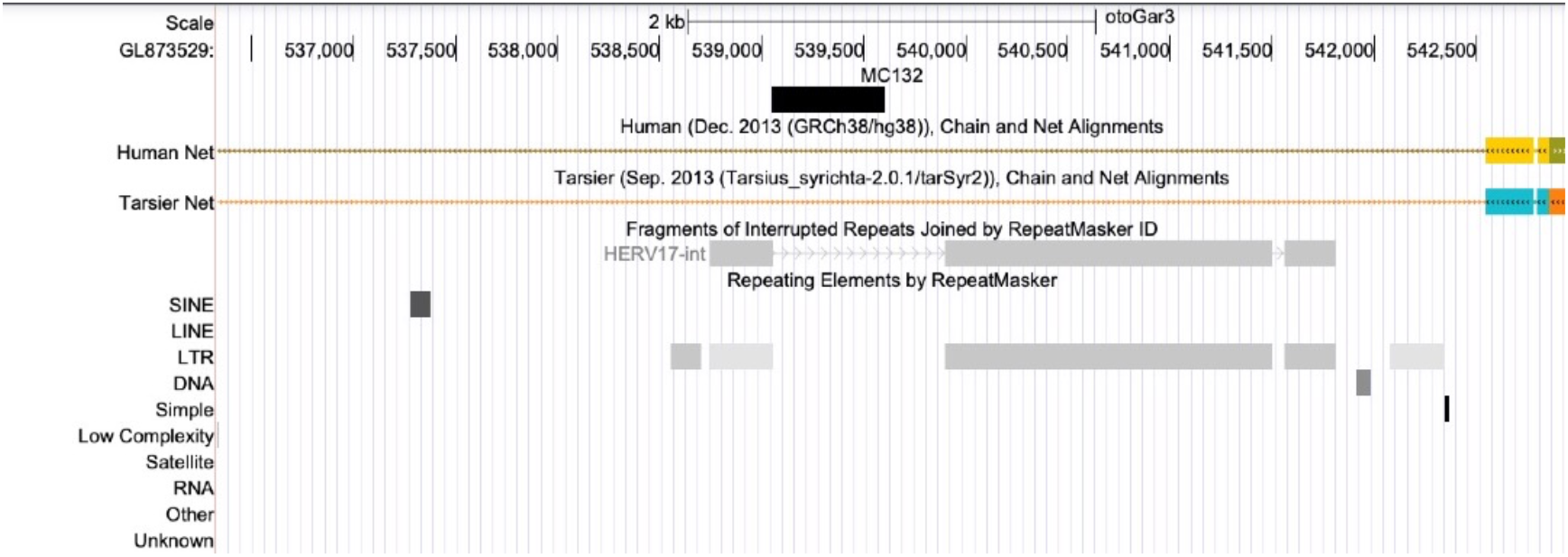
The new protein matches a gap in ERV annotation in the bushbaby genome. The black bar shows a genome segment homologous to MC132, and the gray bars show RepeatMasker annotations. The colored bars show alignments to human and tarsier genomes: these alignments do not cover the segment homologous to MC132.

## DISCUSSION

Some primate retroviruses used to encode an additional protein, homologous to an immune-suppressing protein in a human poxvirus. It is plausible that this retrovirus protein also had an immune-suppressing function. Perhaps some extant retroviruses still encode such a protein. We found one intact ORF for this protein in human chromosome 4, so it is possible that this protein is present in humans. The ORF might have been co-opted as a gene that (down-)regulates immune responses.

One puzzle is that the ORF upstream of gag in a retrovirus would be expected to hinder translation of gag. So it is possible that the ORF’s stop codon inferred by Refiner is incorrect, and it was actually one long ORF fused to gag.

Since retroviruses and poxviruses are not closely related, DNA encoding this protein must have been horizontally transferred between these types of virus. The direction of transfer is unknown, and could be indirect, e.g. from an unknown third source. In any case, a retrovirus encoding this protein infected ancestral primates. It is not clear whether the retrovirus was endogenous or exogenous when it first acquired the protein. Perhaps it infected an ancestor of all primates (Strepsirrhini and Haplorhini), leaving a few traces in bushbaby (and possibly in not-yet-sequenced DNA of lemurs or tarsiers), before proliferating in monkeys. Alternatively, this retrovirus may have independently infected ancestors of monkeys and bushbaby. Homologs of this protein may lurk elsewhere, and finding them should clarify its evolutionary history.

## METHODS

### DNA-versus-protein homology search

DNA-versus-protein homology searches were done with LAST version 1411, essentially as described previously [6]:

~~~
lastdb -q -c myDB proteins.fasta
lastal -D1e9 -K1 -p my.train myDB genome.fasta > out.maf
~~~

This requires a file “my.train” specifying rates of substitution, deletion, and insertion. These rates can be inferred by finding homologies between DNA and protein sequences using last-train [5,6]. However, it is not obvious which sequences to use for this inference: they must have extensive-enough homology to infer the parameters of a 64–21 substitution matrix. We used human pseudogene DNA and non-human proteins: the idea is that they have diverged by a combination of protein-coding and noncoding evolution. Specifically, we used retrogene DNA in human genome hg38 according to ucscRetroInfo9 [13], and chicken proteins from UniProt release 2022_03 (proteome UP000000539) [14]. Instead of chicken, we also tried mouse and zebrafish, but it did not seem to make much difference. The rates were inferred like this:

~~~
lastdb -q -c myChickenDB UP000000539_9031.fasta
last-train --codon --pid=50 -m100 myChickenDB pseudos.fasta > my.train
~~~

### DNA consensus sequences

Refiner, from RepeatModeler version 2.0.3, was run with default parameters.

## ACKNOWLEDGEMENTS

We are grateful to Junna Kawasaki, Atsushi Takeda, and Michiaki Hamada for discussions about viral fossils, and to Wojciech Makałowski for comments on “homology”.

## REFERENCES

[1] Wang J, Han GZ. Frequent retroviral gene co-option during the evolution of vertebrates. Molecular Biology and Evolution. 2020 Nov;37(11):3232–42.

[2] Thompson PJ, Macfarlan TS, Lorincz MC. Long terminal repeats: from parasitic elements to building blocks of the transcriptional regulatory repertoire. Molecular cell. 2016 Jun 2;62(5):766–76.

[3] Johnson WE. Origins and evolutionary consequences of ancient endogenous retroviruses. Nature Reviews Microbiology. 2019 Jun;17(6):355–70.

[4] Brady G, Haas DA, Farrell PJ, Pichlmair A, Bowie AG. Poxvirus protein MC132 from molluscum contagiosum virus inhibits NF-κB activation by targeting p65 for degradation. Journal of Virology. 2015 Aug 1;89(16):8406–15.

[5] Yao Y, Frith M. Improved DNA-versus-Protein Homology Search for Protein Fossils. IEEE/ACM Transactions on Computational Biology and Bioinformatics. 2022 May 26.

[6] Frith MC. Paleozoic Protein Fossils Illuminate the Evolution of Vertebrate Genomes and Transposable Elements. Molecular biology and evolution. 2022;39(4):msac068.

[7] Storer J, Hubley R, Rosen J, Wheeler TJ, Smit AF. The Dfam community resource of transposable element families, sequence models, and genome annotations. Mobile DNA. 2021 Dec;12(1):1–4.

[8] Hubley R, Wheeler TJ, Smit AF. Accuracy of multiple sequence alignment methods in the reconstruction of transposable element families. NAR genomics and bioinformatics. 2022 Jun;4(2):lqac040.

[9] Kent WJ, Sugnet CW, Furey TS, Roskin KM, Pringle TH, Zahler AM, Haussler D. The human genome browser at UCSC. Genome research. 2002 Jun 1;12(6):996–1006.

[10] Grandi N, Cadeddu M, Blomberg J, Tramontano E. Contribution of type W human endogenous retroviruses to the human genome: characterization of HERV-W proviral insertions and processed pseudogenes. Retrovirology. 2016 Dec;13(1):1–25.

[11] Pavlíček A, Pačes J, Elleder D, Hejnar J. Processed pseudogenes of human endogenous retroviruses generated by LINEs: their integration, stability, and distribution. Genome research. 2002 Mar 1;12(3):391–9.

[12] Mi S, Lee X, Li XP, Veldman GM, Finnerty H, Racie L, LaVallie E, Tang XY, Edouard P, Howes S, Keith JC. Syncytin is a captive retroviral envelope protein involved in human placental morphogenesis. Nature. 2000 Feb;403(6771):785–9.

[13] Baertsch R, Diekhans M, Kent WJ, Haussler D, Brosius J. Retrocopy contributions to the evolution of the human genome. BMC genomics. 2008 Dec;9(1):1–9.

[14] The UniProt Consortium. UniProt: the universal protein knowledgebase in 2021. Nucleic Acids Res. 49:D1.

